# Circulating miR-29a as a new biomarker of food anaphylaxis and endothelial glycocalyx regulation

**DOI:** 10.64898/2026.04.06.716635

**Authors:** Antonio Muñoz-Callejas, Adrián Moreno-Vidal, Alejandro Henar-Izquierdo, Lucía Palacio-García, Sergio Fernández-Bravo, Irene de María-Camacho, Angela Di Giannatale, Alicia Gómez-López, Pablo Rodríguez Del Rio, José Julio Laguna, Alberto Benito-Martin, Emilio Nuñez-Borque, Vanesa Esteban

## Abstract

**Background:** Anaphylaxis is an acute and potentially life-threatening hypersensitivity reaction often involving the cardiovascular system. Circulating microRNAs (miRNAs/miR), including those carried by extracellular vesicles (EVs), are emerging biomarkers that display regulatory functions in allergy. This study aims to investigate the role of miR-29a in anaphylaxis.

**Methods:** MiR-29a (3p and 5p) levels were assessed by qPCR from acute and baseline samples of serum and EVs from 70 patients with food- and drug-mediated anaphylaxis. EVs purification was confirmed by Western blot, electron microscopy, and NanoSight. MiR-29a-3p target genes were studied *in silico* using systems biology analysis (SBA). Moreover, miR-29a levels were evaluated *in vitro* in endothelial cells (ECs) exposed to anaphylactic mediators. Additionally, a panel of endothelial glycocalyx (eGCX)-associated mRNA was analyzed after transfection with a miR-29a-3p inhibitor.

**Results:** Patients with food-induced anaphylaxis exhibited reduced miR-29a-3p levels in both serum and EVs during the acute reaction. In contrast, miR-29a-5p levels were decreased in serum but not in EVs. No significant modulation of either miRNA was observed in drug-induced anaphylaxis. SBA of miR-29a-3p identified molecular pathways, biological processes and functional networks associated with eGCX remodelling. Intracellular levels of miR-29a-3p were modulated *in vitro* in ECs following exposure to anaphylactic mediators. Inhibition of miR-29a-3p significantly reduced *ESM1* expression.

**Conclusions:** The miR-29a-3p levels are decreased in serum and EVs from patients with acute food-induced anaphylaxis, suggesting its potential as a promising biomarker. Moreover, a role for miR-29a-3p in eGCX integrity under anaphylactic conditions was demonstrated, potentially regulating *ESM1*.

**Key Message:** MiR-29a-3p is selectively reduced in serum and extracellular vesicles during acute food-induced anaphylaxis and may regulate endothelial glycocalyx-related pathways, which supports its potential as a novel biomarker and molecular mediator of vascular involvement in anaphylactic reactions.

## Introduction

Anaphylaxis is a severe form of allergic reaction and may be potentially life-threatening^1^. Several allergens can trigger these reactions including food, drugs, and Hymenoptera venoms. Specifically, drugs represent the most common cause in adults, while food is the predominant trigger in children^2,3^. During an anaphylactic reaction, multiple symptoms converge into a systemic response where mucous membranes and skin are the organs most frequently involved^4^. Notwithstanding, the most severe cases are determined by hypoxia and/or hypotension^5^. Therefore, the cardiovascular system is among the most severely affected during anaphylactic shock and, among its components, the endothelium is particularly altered. This barrier consists of a monolayer of endothelial cells (ECs) and participates in numerous vital functions^6^, playing a key role in the vascular permeability associated with anaphylaxis^7,8^. Moreover, ECs present a dynamic structure on their surface called endothelial glycocalyx (eGCX), mainly composed of glycoproteins and proteoglycans. This malleable sugar-based structure is essential for communication between tissues and the bloodstream^9^, shifting accordingly to the haemostatic conditions and playing a pleiotropic role in health and disease^10^. The plasticity of the eGCX makes it highly prone to damage, exposing the endothelial membrane and contributing to endothelial dysfunction, as seen in pro-inflammatory diseases such as sepsis or atherosclerosis^11,12^. Precisely, eGCX is quickly damaged in mice models of anaphylaxis, causing the release of the endothelial cell specific molecule-1 (Endocan 1; *ESM1*)^13^.

Mechanistically, anaphylaxis is mainly triggered by the IgE pathway, which gives rise to the release of several mediators such as tryptase, histamine, or the platelet-activating factor (PAF). These molecules cause the signs and symptoms of this pathological event^14^. However, IgE-independent pathways have also been described^15^. In recent years, other mediators, such as extracellular vesicles (EVs) and microRNAs (miRNAs or miR)^4^, have also been described in anaphylaxis, demonstrating the molecular complexity of these reactions. Although some endotypes and phenotypes have been recognized in anaphylaxis^2,16^, their translation into precision medicine strategies for patients management remains limited.

The diagnosis of anaphylaxis relies on clinical signs and symptoms, which overlap with other pathologies. Moreover, there is an absence of reliable diagnostic markers to confirm the diagnosis of anaphylaxis. In this regard, serum tryptase, the main biomarker for mast cell related responses, provides limited utility because it is not always increased in anaphylactic patients^17^. Therefore, it is necessary to find novel molecules capable of improving the diagnosis and management of patients with anaphylaxis.

MiRNAs are small sequences of non-coding RNA of approximately 22 nucleotides, whose main function is the regulation of mRNA translation. Therefore, they play a key role in post-transcriptional gene control and in maintaining the homeostasis^18^. Moreover, miRNAs are easy to obtain and stable molecules, both desirable features of a good biomarker^19^. Our group demonstrated that children with food-induced anaphylactic reactions present high levels of miR-21-3p and miR-487b-3p^20^. In addition, we have also described lower serum levels of miR-375-3p in patients with drug- and food-induced anaphylaxis^21^. In parallel, Francuzik *et al*. identified high serum levels of the miR-451a in adults with anaphylaxis by food and Hymenoptera venom^22^.

MiRNAs can circulate in different ways: free, complexed with other proteins, or within EVs^23^. Interestingly, miRNAs transported by EVs are protected from degradation by nucleases, allowing them to reach different and distant cellular niches^24^. Our group described an increase in miR-21-3p and a decrease in miR-375-3p levels within EVs during anaphylaxis^21,24^. Moreover, functional studies of these miRNAs suggest that they may participate in the immunological and vascular mechanisms of the anaphylactic reaction^24^. Precisely, EVs from patients with anaphylaxis can induce an increase in endothelial permeability^25^.

Currently, different miRNAs have been associated with cardiovascular and allergic diseases^26,27^. Among them, the miR-29 has been described as an important regulator of different functions such as proliferation, fibrosis, and immune response^28^. In allergic diseases, it has been found to be elevated in nasal samples from patients with rhinitis, where it compromises epithelial barrier integrity^29^. On the other hand, reduced plasma levels of this miRNA have been linked to vascular endothelial damage^28^, and its inhibition impairs vasodilation^30^. In addition, it has been reported to play a protective role by decreasing the expression of adhesion molecules and reducing endothelial dysfunction both *in vitro* and *in vivo* models^31^. Nevertheless, despite its known role in vascular damage and in allergic diseases, this miRNA has not yet been investigated in the context of anaphylaxis.

Therefore, we aim to characterize the circulating levels of the miR-29a in patients with anaphylaxis by different triggers, and to determine its impact on the endothelium, focusing on eGCX integrity.

## Methods

A supplementary version of the methodology is available on **Supplementary Material.**

### Subjects of study

The study was approved by the Ethics Committee (CEIm FJD, PIC057-19 and PIC166-22). A total of 91 patients with anaphylaxis were recruited from different Spanish hospitals. However, 21 adult patients were removed due to the hemolysis present in one, or both, of their samples. Finally, a total of 70 patients, 51 adults (>18 years old) and 19 children (<18 years old) were ultimately included in the study. Diagnosis was determined by an allergist based on the criteria established by the National Institute of Allergy and Infectious Disease and Food Allergy and Anaphylaxis Network^32^.

Clinical characteristics of each subject were recorded: age, sex, suspected allergen, presenting symptoms and treatment administration before acute sample collection. Severity was determined according to the criteria established by Brown classification^5^. Serum tryptase levels were measured using UniCAP (Thermo Fisher Scientific). A complete description of patients and their reactions can be found in the **Supplemental Table 1**.

### Sample collection

Serum samples were collected from each patient under two conditions: during the acute phase (anaphylaxis) and at baseline (control), at least 14 days after the reaction. Considering the heterogeneity among subjects, acute phase data were expressed as fold-changes relative to individual baseline values (ratio acute/basal). Paired samples from each patient were always analyzed simultaneously and in the same batch.

### Circulating serum miRNAs analysis

RNA was extracted from 200 µl of serum using the miRNeasy Serum/Plasma Advanced kit (Qiagen) and retrotranscribed by the miRCURY LNA RT kit (Qiagen), according to Nuñez-Borque *et al.*^33^. MiRNAs quantification was performed by quantitative PCR (qPCR) in the LightCycle96 Real Time PCR System (Roche Life Science) using the miRCURY LNA SYBR Green PCR kit (Qiagen) with specific primers (Qiagen; **Supplemental Table 2)**. UniSp2/4/5 and UniSp6 were used as exogenous controls of the extraction and reverse transcription, respectively; miR-451a and miR-23a-3p were measured as endogenous controls of serum hemolysis^34,35^; and miR-30e-5p was chosen as endogenous control for the normalization of the target miRNAs (miR-29a-3p and miR-29a-5p), as previously used in other studies^20,21^. Data obtained were analyzed using the 2^-ΔΔCT^ method^36^.

### Determination of miRNAs within extracellular vesicles

The purification of EVs was carried out from 500 µl of serum using the miRCURY Exosome isolation Kit – Serum and Plasma (Qiagen), according to Colletti *et al*.^37^. Suitable isolation and characterization of EVs was determined by Western blot (WB), electron microscopy (EM), and nanoparticle tracking analysis (NTA), following MISEV2023 guidelines^38^. The extraction, reverse transcription, and quantification of miRNAs contained in EVs were carried out as described in Nuñez-Borque *et al.*^33^.

### Systems biology analysis

Systems Biology Analysis (SBA) was performed *in silico* using all miR-29a-3p target genes with ≥50 target score in the miRDB database (http://mirdb.org/). All genes (1015 genes) were uploaded to ShinyGO 8.0 (https://bioinformatics.sdstate.edu/go/) and to the search tool for retrieving interacting genes or proteins (STRING; https://string-db.org/) for analysis.

### Determination of miR-29a levels in stimulated ECs

Human aortic ECs (HAEC; Lonza CC-2535) and Human dermal microvascular ECs (HMVEC-D; Lonza CC-2543) were grown and seeded until monolayer formation (90% confluence) in plates pretreated with 0.5% gelatin (Sigma-Aldrich).

Before the stimulation, HAEC and HMVEC-D were serum bovine fetal (FBS) depleted (0.5%) for 24 hours. After this time, 0.1 μM of PAF (Sigma-Aldrich) and 1 μM of Histamine (Sigma-Aldrich) were added and incubated for 2 hours at 37°C in 5% CO□ (EC-anaphylaxis). Medium was used as a negative control (EC-control). Extraction of miRNAs was carried out using the Master Pure Complete DNA & RNA Purification Kit (Lucigen). Reverse transcription and miRNAs quantification were performed as described in the circulating serum miRNAs section.

### Transfection of miRNA and gene expression analysis in HAEC

HAECs were seeded and grown in 6-well plates (Corning Costar) for 24 hours to 70-80% confluence. Transfection was performed by incubating cells with Opti-MEM medium (Thermo Scientific) and with the Transit-X2 reagent (Qiagen) combined with an inhibitor of the miR-29a-3p at 50 mM (Qiagen). In addition, a fluorescently labeled inhibitor scramble (Qiagen) was used as a control (50 mM) to confirm the successful transfection of the cells by LSM 700 inverted confocal microscope (Carl Zeiss).

For RNA extraction, cell scrapers and 1 ml of TRIzol (Invitrogen) were employed to lysate cells. RNA was then retrotranscribed using High-Capacity cDNA Reverse Transcription kit (Qiagen) and the ProFlex PCR system thermal cycler (Applied Biosystems, Waltham). Quantification of mRNAs was performed using the TB Green Premix Ex Taq (Takara) in the Quant Studio 3 Real-Time PCR System (Thermo Fisher Scientific). The 18s rRNA was used as endogenous control for the normalization of the target mRNAs. Expression of various proteins related to the eGCX was determined using specific primers (Eurofins, **Supplementary Table 3**). Data obtained were analyzed using the 2^-ΔΔCT^ method^36^.

### Statistical analysis

Data analysis and graphical representation were performed using the GraphPad Prism 8.01 Software (La Jolla, CA, USA). Qualitative variables were described as frequency and percentage. Graphs were represented by the mean ± standard error of the mean (SEM) in parametric analyses, while it showed the median ± interquartile range (IQR) in non-parametric tests. Wilcoxon matched-pairs test was used for serum, EVs and ECs data, while Mann-Whitney test was applied for comparison between patients with and without cardiovascular symptoms, and Unpaired t test was used for transfection experiments. Fold-change normalized data were analyzed, establishing the value of 1 to the control condition in each case. Statistical significance was determined using two-tailed tests and p-values were considered significative at level of p<0.05.

## Results

### Characteristics of the patient cohort

The study population initially comprised 182 paired serum samples obtained from 91 patients with anaphylaxis by different triggers. However, after excluding hemolyzed samples, the studied cohort included 70 patients with food (51%) or drug (49%) induced anaphylaxis. The detailed distribution of samples studied is described in **Supplementary Figure 1A.** Women represented the largest proportion (61%) and the age of subjects ranged from 4 to 76 years. The most frequent manifestations were cutaneous (84%) and respiratory (89%) ones, followed by digestive (61%) and mucosal (50%) symptoms, while cardiovascular (31%) and nervous (19%) systems were the least commonly affected. Accordingly, 68% of reactions were classified as moderate (Grade 2) and 31% as severe (Grade 3). Notably, severe reactions were more frequent in adults (35%) than in children (10%). Moreover, 75% of patients received treatment prior to sample collection, strictly prioritizing clinical care. Among treated individuals before collecting acute sample, the most frequently administered treatments were corticosteroids (91%), H1R antagonists (89%), and epinephrine (77%), whereas β2-adrenergic agonists (24%) and H2R antagonists (13%) were less commonly used (**Table 1**).

**Table 1.** Clinical characteristics of patients and features of their hypersensitivity reactions. * Severity was graded according to Brown classification^5^. ^#^ Percentage of patients treated prior to acute sample collection; remaining subjects received treatment after obtaining the acute phase serum. Significant differences between groups (p<0.05) are indicated in **bold**.

|  | Patients |  |  |  | p-value |  |  |
| --- | --- | --- | --- | --- | --- | --- | --- |
|  | Total | AD | AF | CF | AD vs AF | AD vs CF | AF vs CF |
| **Number** | 70 | 34 | 17 | 19 | NA | NA | NA |
| **Sex (female; %)** | 61.4 | 64.7 | 58.8 | 57.8 | 0.882 | 0.631 | 0.789 |
| **Age (years) ± SEM** | 34.18 ± 19 | 46.44 ± 14 | 35.23 ± 15 | 11.31 ± 4 | **0.013** | **<0.0001** | **<0.0001** |
| **Severity (%) \*** |  |  |  |  |  |  |  |
| **Grade 2** | 61.4 | 47.0 | 64.7 | 84.2 | 0.242 | **0.007** | 0.187 |
| **Grade 3** | 38.5 | 52.9 | 35.2 | 15.7 | 0.242 | **0.007** | 0.187 |
| **Symptoms (%)** |  |  |  |  |  |  |  |
| **Skin** | 84.2 | 88.2 | 94.1 | 68.4 | 0.515 | 0.079 | 0.053 |
| **Mucous** | 50.0 | 41.1 | 70.5 | 47.3 | 0.209 | 0.609 | 0.505 |
| **Digestive** | 61.4 | 55.8 | 76.4 | 57.8 | 0.157 | 0.889 | 0.250 |
| **Respiratory** | 88.5 | 91.1 | 76.4 | 94.7 | 0.156 | 0.645 | 0.120 |
| **Nervous** | 18.5 | 32.3 | 0.0 | 10.5 | **0.007** | 0.079 | 0.178 |
| **Vascular** | 31.4 | 38.2 | 29.4 | 21.0 | 0.543 | 0.206 | 0.576 |
| **Treatment (%)** |  |  |  |  |  |  |  |
| **Before collecting sample #** | 75.7 | 85.2 | 82.3 | 52.6 | 0.790 | **0.009** | 0.061 |
| Epinephrine | 57.1 | 64.7 | 52.9 | 47.3 | 0.427 | 0.227 | 0.747 |
| H1R antagonist | 54.2 | 55.8 | 70.5 | 36.8 | 0.320 | 0.190 | **0.044** |
| H2R antagonist | 10.0 | 8.8 | 23.5 | 0.0 | 0.156 | 0.189 | **0.024** |
| Corticosteroids | 57.7 | 64.7 | 6.8 | 26.3 | **0.004** | **0.006** | 0.852 |
| β2-adrenergic agonist | 14.7 | 14.7 | 11.7 | 21.0 | 0.116 | 0.390 | 0.469 |

AD: adults with drug-induced anaphylaxis; AF: adults with food-induced anaphylaxis; CF: children with food-induced anaphylaxis; NA: not applicable; H1R: histamine receptor 1; H2R: histamine receptor 2.

### Serum miR-29a levels are reduced in patients with food-mediated anaphylaxis

To determine changes in miR-29a levels during anaphylaxis, serum samples from 74 patients with food- and drug-mediated reactions were analyzed. However, 14 patients were excluded from the analysis due to hemolysis in one or both samples (**Figure 1A, Supplementary Figure 1B)**. The stepwise detection of UniSp2/4/5 (**Figure 1B**) and the homogeneous detection of UniSp6 (**Figure 1C**) among samples confirmed the correct extraction and reverse transcription of miRNAs in the remaining 60 patients.

**Figure 1.**
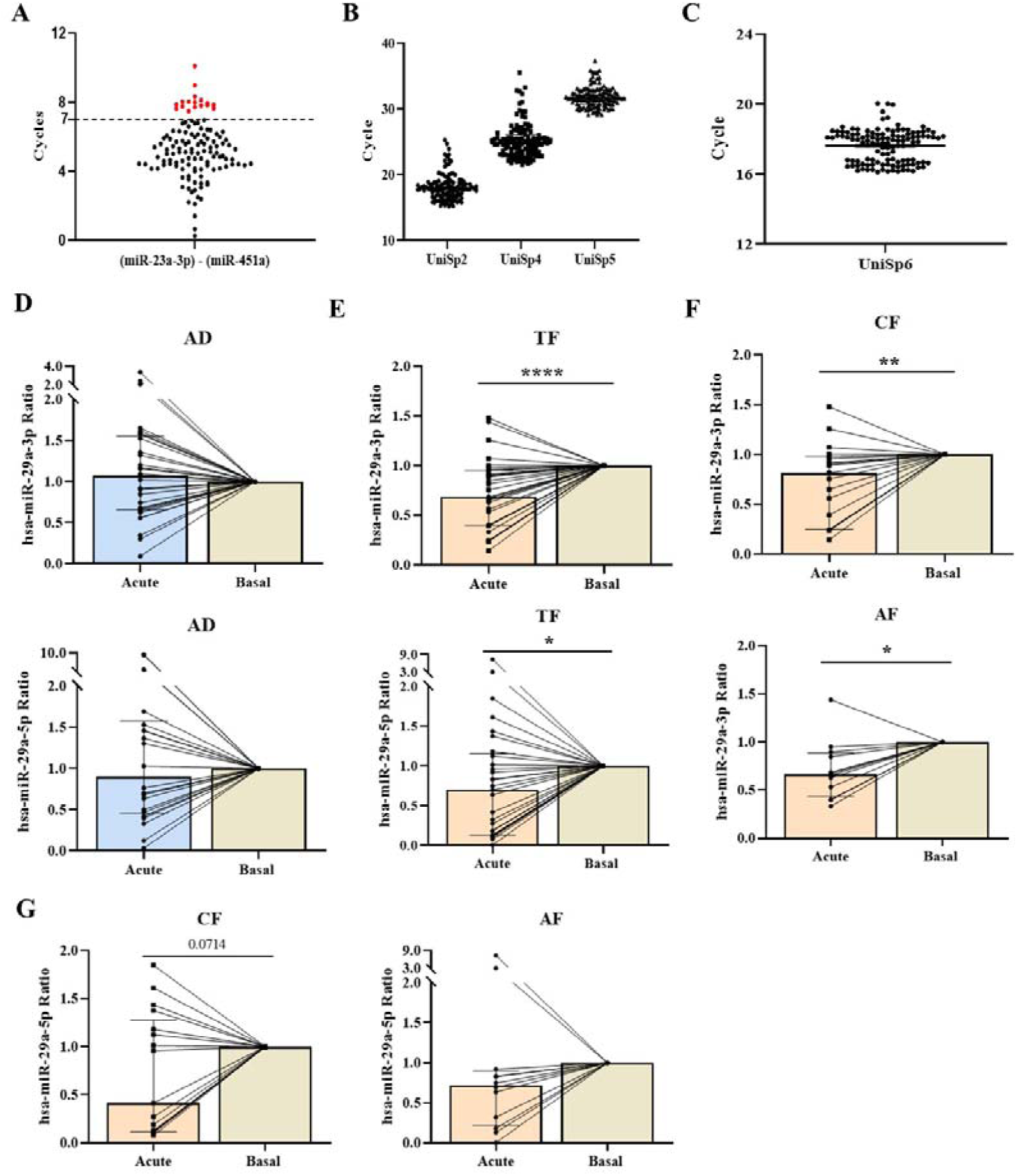
Determination of miR-29a serum levels in patients with anaphylaxis. (**A**) Sample quality was assessed by the ratio between miR-23a-3p and miR-451a (≥ 7 cycles). 14 patients were discarded from the study due to the hemolysis in one, or both, of their samples (red points). (**B**) Quality of miRNA extraction was verified using the exogenous controls UniSp2, UniSp4, and UniSp5. (**C**) The correct development of reverse transcription was confirmed by measuring the exogenous control UniSp6. (**D**) Serum levels of miR-29a-3p (n=29, up) and miR-29a-5p (n=22, down) in adults with drug-induced anaphylaxis (AD). (**E**) Serum levels of miR-29a-3p (n=31, up) and miR-29a-5p (n=29, down) in all patients with food-induced anaphylaxis (TF). (**F-G**) Serum levels of miR-29a-3p (**F**) and miR-29a-5p (**G**) stratifying between adults (AF; n=12) and children (CF; n=19) with food-induced anaphylaxis. Each point represents the ratio between the acute phase and the baseline from each patient. Differences in the number of patients analyzed are due to the non-detection of miRNA in any of the samples from those patients. Data show the median ± IQR. Statistical analyses were performed using Wilcoxon test. *p<0.05, **p<0.01, ****p<0.0001.

Once the quality of the samples and the techniques were confirmed, the levels of miR-29a-3p and miR-29a-5p were measured in the study cohort. The analysis performed in patients with drug-induced anaphylaxis revealed no differences between the acute and basal conditions (**Figure 1D**). In contrast, a decrease during the reaction of miR-29a-3p and miR-29a-5p was observed in patients with food-induced anaphylaxis (**Figure 1E**). However, when stratified between adults and children, these differences were only observed in the case of miR-29a-3p (**Figure 1F**), but not in miR-29a-5p (**Figure 1G**).

### MiR-29a levels decrease within the EVs of patients with food-mediated anaphylaxis

Considering the relevance of miRNAs transported by EVs as long-distance mediators, we decided to determine if the reduction in miR-29a-3p and miR-29a-5p levels observed in serum also occurred within the EVs of patients with anaphylaxis. The correct purification of these particles was first confirmed by immunodetection of specific markers (**Figure 2A**), by their visualization using EM (**Figure 2B**), and by their characterization through NTA (**Figure 2C**), where a heterogeneous population of EVs was observed.

**Figure 2.**
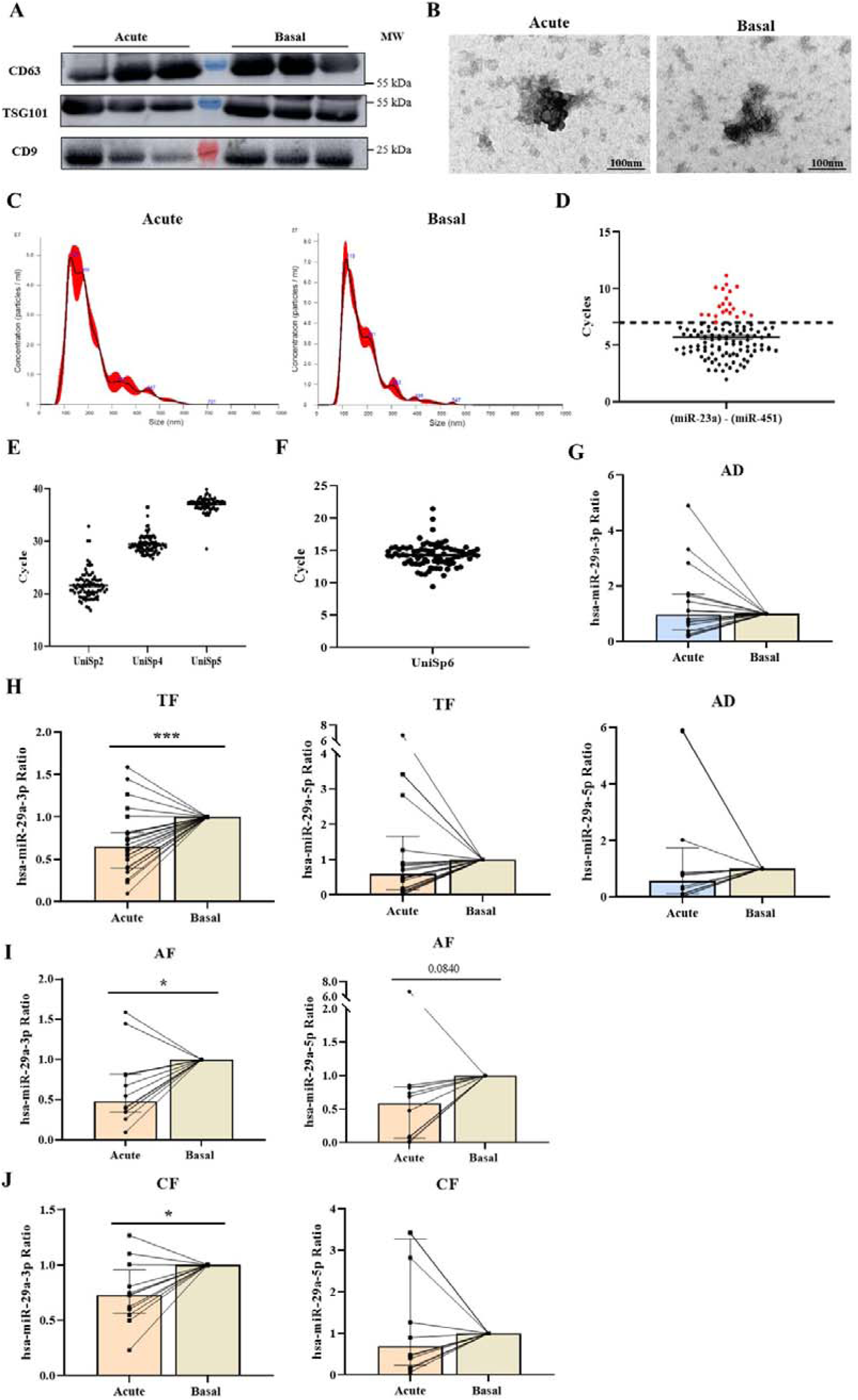
Analysis of miR-29a levels within EVs from patients with anaphylaxis. (**A-C**) Characterization of EVs purified from serum of patients with anaphylaxis during the acute and basal phase: (**A**) detection of EVs proteins (CD63, TSG101 and CD9) by Western blot, (**B**) visualization of EVs by electron microscopy (Scale: 100 nm), and (**C**) characterization of particle size (nm) and concentration (particles/ml) by NanoSight. Acute and basal samples from three adults with food-induced anaphylaxis were analyzed. (**D**) Sample quality was assessed by the ratio between miR-23a-3p and miR-451a. 16 patients were discarded from the study due to exceeding the established threshold (≥7 cycles) in one, or both, of their samples (red points). (**E**) The quality of miRNA extraction was assessed using UniSp2, UniSp4, and UniSp5. **(F)** The efficiency of reverse transcription was verified by quantifying the UniSp6. (**G**) Levels of miR-29a-3p (n=16, up) and miR-29a-5p (n=12, down) within EVs of adults with drug-induced anaphylaxis (AD). (**H**) Levels of miR-29a-3p (n=24, left) and miR-29a-5p (n=22, right) within EVs of all patients with food-induced anaphylaxis (TF). (**I**) Levels of miR-29a-3p (n=12, left) and miR-29a-5p (n=10, right) within EVs of adults with food-induced anaphylaxis (AF). (**J**) Levels of miR-29a-3p (n=12, left) and miR-29a-5p (n=12, right) within EVs of children with food-induced anaphylaxis (CF). Each point represents the ratio between the acute phase and the baseline from each patient. Differences in the number of patients analyzed are due to the non-detection of miRNA in any of the samples from those patients. Data show the median ± IQR. Statistical analyses were performed using Wilcoxon test. *p<0.05***, p<0.001; MW: molecular weight.

EVs from 56 patients with anaphylaxis were purified to measure the miR-29a levels. However, 16 subjects were discarded due to the hemolysis of their samples (**Figure 2D, Supplementary Figure 1C**). The correct performance of the techniques was checked using the UniSp2/4/5 (**Figure 2E**) and the UniSp6 (**Figure 2F**) in the remaining 40 patients.

Similarly to the results obtained in serum, the specific analysis of miR-29a levels showed no changes in adults with drug-induced anaphylaxis (**Figure 2G**). However, in patients with food-induced anaphylaxis, miR-29a-3p levels were significantly reduced during the reaction, whereas no changes were observed in miR-29a-5p (**Figure 2H**). Notably, both adults (**Figure 2I**) and children (**Figure 2J**) cohorts showed decreased levels of miR-29a-3p.

### Intracellular levels of miR-29a-3p modulate eGCX integrity in ECs

When serum miR-29a levels were analyzed accordingly to the specific trigger of the reaction, to the treatment received before collecting the acute sample, or to sex, no differences were observed (data not shown). However, miR-29a-3p levels were markedly decreased in patients with food-induced anaphylaxis presenting cardiovascular symptoms (**Figure 3A**). Considering the endothelium as the main structure affected in cardiovascular alterations underlying anaphylaxis^7,14^, we evaluated the modulation of miR-29a intracellular levels in macrovascular (HAEC) and microvascular (HMVEC-D) ECs under stimulation with key mediators of anaphylaxis: histamine and PAF. Similarly to the results in serum and EVs, no variations in intracellular miR-29a-5p levels were observed in ECs after incubation with anaphylactic mediators (**Figure 3B**). Instead, miR-29a-3p levels intracellularly increased in HAEC and decreased in HMVEC-D after the stimulation (**Figure 3C**), suggesting a different role for this miRNA depending on the endothelial niche.

**Figure 3.**
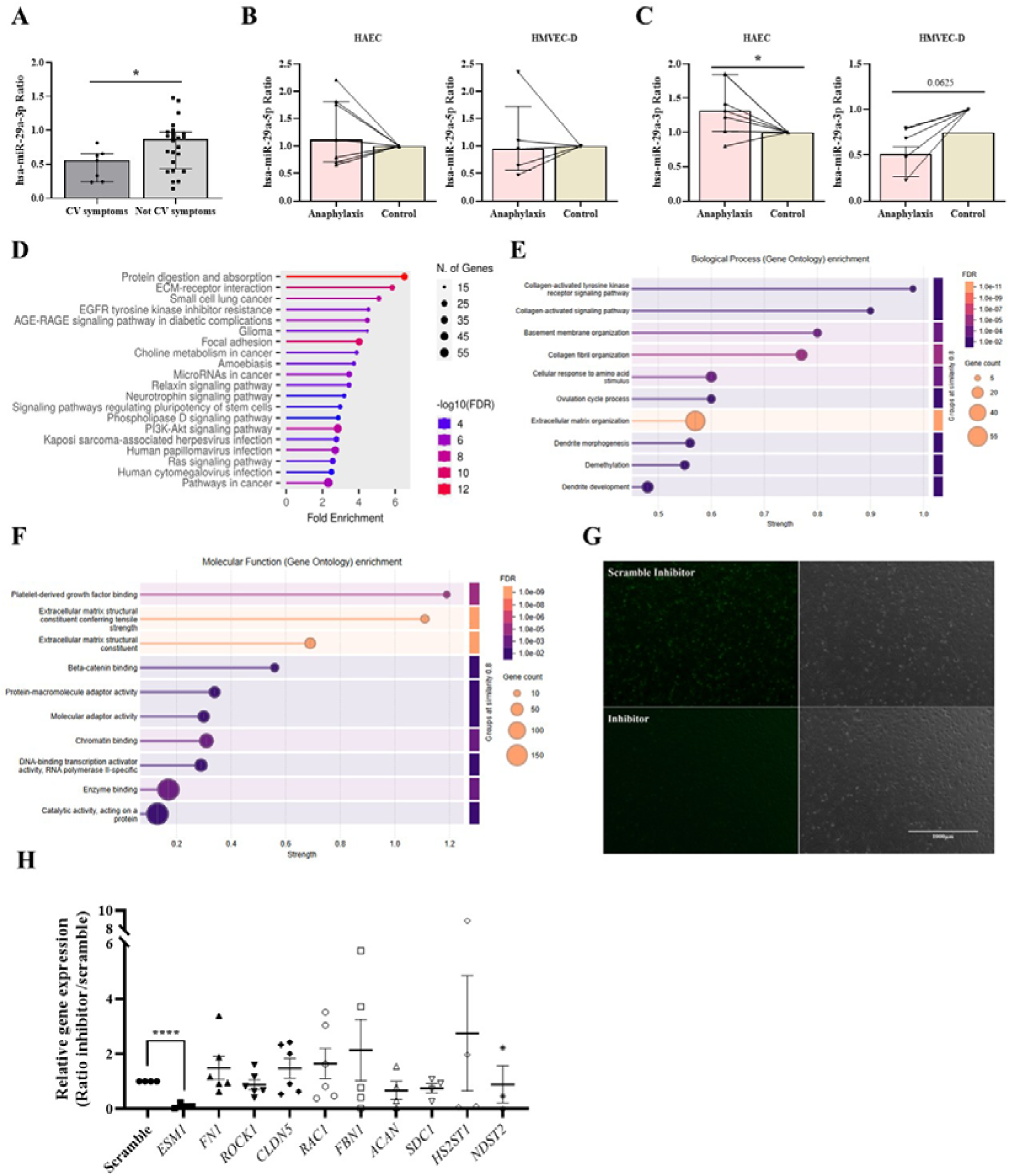
Evaluation of the involvement of miR-29a-3p in the endothelium during anaphylaxis. (**A**) Comparison of serum miR-29a-3p levels between patients with cardiovascular (CV) symptoms (n=7) and those without them (n=24) in all patients with food-induced anaphylaxis. Data show the median ± IQR. Statistical analyses were performed using Mann-Whitney test. (**B-C**) Quantification of intracellular miR-29a-5p (**B**) and miR-29a-3p (**C**) levels in human aortic endothelial cells (HAECs; n=7) and human dermal microvascular endothelial cells (HMVEC-D; n=5) after 2 h of stimulation with anaphylactic mediators (histamine and platelet activating factor). Each point represents the ratio between stimulated and non-stimulated conditions. Data show the median ± IQR. Statistical analyses were performed using Wilcoxon test. (**D-F**) *In silico* analysis of miR-29a-3p target genes. Most significant ‘Pathways’ (**D**), obtained by ShinyGO, and most significant ‘Biological Process’ (**E**) and ‘Molecular Function’ (**F**), obtained by STRING. (**G**) Representative images of HAECs transfected with miR-29a-3p inhibitor and its fluorescent scramble control (green). The left panels show fluorescence images, while right panels show brightfield images. Scale bar: 1000 µm. All images were obtained using fluorescence microscopy with a 20x objective. (**H**) Levels of the different endothelial glycocalyx (eGCX) genes after the transfection with the miR-29a-3p inhibitor (n=3-6): Endocan 1 (*ESM1)*, Fibronectin 1 (*FN1)*, Rho-associated protein kinase 1 (*ROCK1)*, Claudin 5 (*CLDN5)*, Rac family small GTPase 1 (*RAC1)*, Fibrillin 1 (*FBN1)*, Aggrecan (*ACAN)*, Syndecan 1 *(SDC)*, Heparan sulfate 2-O-sulfotransferase 1 (*HS2ST1)*, N-Deacetylase and N-Sulfotransferase 2 (*NDST2)*. Each point represents the ratio between the miR-29a-3p inhibitor and scramble. Data show the mean ± SEM. Statistical analyses were performed using unpaired t test. *p<0.05, ****p<0.0001; FDR: false discovery ratio.

To investigate functional implications of these changes in the endothelium, we performed an *in silico* analysis focusing on miR-29a-3p target genes. SBA showed a stronger relationship with pathways involved in the extracellular matrix organization and maintenance (**Figure 3D-F**). Precisely, miR-29a has been described to target different extracellular matrix components, such as collagen, elastin and fibrillin 1^39^. Therefore, changes in endothelial miR-29a-3p could modulate eGCX, a component of the extracellular matrix whose alteration has been described in anaphylaxis^13^. To investigate this, we performed an *in vitro* inhibition of miR-29a-3p and analyzed its impact on different eGCX components in HAEC. Immunofluorescence images confirmed successful transfection of the miR-29a-3p (**Figure 3G**). However, its blockade did not show significant changes in the levels of most of the genes analyzed. In contrast, the expression of *ESM1* was significantly reduced (**Figure 3H**), suggesting an impact of miR-29a-3p on eGCX remodelling.

## Discussion

The interest in miRNAs has increased in recent years, as they represent novel molecular mediators and promising biomarkers in the field of allergy^26^. In this work, we found that miR-29a levels decreased in acute samples (serum and EVs) of patients with food-induced anaphylaxis, while no changes were seen in those with drug-mediated reactions, indicating that variations on this miRNA could be specific to anaphylaxis caused by food. The miR-29a-3p is necessary for the proper functioning of the endothelium^30^; thus, we evaluated *in vitro* the changes occurring in the macro- and microvasculature, demonstrating that miR-29a-3p levels increased in HAEC and decreased in HMVEC-D under anaphylactic stimuli. To determine the mechanistic effect of these variations on ECs, a SBA of their target genes was performed and different signaling pathways were examined, among which the stability of the extracellular matrix stood out. Therefore, since eGCX is a relevant component of the extracellular matrix in the macrovascular system^40^, we inhibited miR-29a-3p in HAEC and observed a reduction in the expression of *ESM1*, which has been recently linked to anaphylaxis^13^.

In anaphylaxis, only three studies have characterized serum miRNA profiles in humans: i) one in adults with drug-induced anaphylaxis, which validated a decrease in miR-375-3p^20^; ii) one in children with food-mediated anaphylaxis, which validated an increase in miR-21-3p and miR-487b-3p^21^; iii) and another in adults with food- and Hymenoptera venom-induced anaphylaxis, which validated an increase in miR-451a^22^. Our findings demonstrated a decrease in serum miR-29a levels in patients with food-mediated anaphylaxis, but not in those with drug-mediated anaphylaxis. Changes in this miRNA did not reach statistical significance in previous next generation sequencing studies, possibly due to the small sample sizes used in these analyses and/or the heterogeneity among anaphylactic reactions. However, our data show a clear difference in miR-29a levels depending on the trigger of the reaction, demonstrating endotypic differences based on the type of allergen, and suggesting that it could be a potential biomarker for distinguishing atopic reactions.

EVs-associated miRNAs are particularly relevant as mediators of intercellular communication, since vesicular transport protects them from degradation and allows miRNAs to reach distant target sites^24^. This mechanism is especially important in a systemic reaction such as anaphylaxis. However, although a characteristic protein profile has been described in EV from patients with anaphylaxis^25^, few details are known about the relevance of these particles and their content in the development of the anaphylactic reaction. In this study, we found that miR-29a-3p levels were reduced in EVs from patients with food-induced anaphylaxis, as we had previously described for miR-375-3p and for the increase of miR-21-3p^21,24^. Currently, it has been demonstrated that miR-29a within EVs contributes to the pathogenesis of other diseases, such as systemic lupus erythematosus^41^. In turn, the overexpression of miR-29a in mesenchymal stem cell exosomes decreases the migration and the blood vessel formation in glioma cells^42^, while in bone stem cells it diminishes fibrosis by reducing extracellular proteins such as collagen I^43^. Moreover, exosomal miR-29a regulates microvascular EC proliferation, migration, and angiogenesis by targeting VEGFA^44^. Therefore, miR-29a-3p carried by EVs could play a role as a mediator in the molecular basis underlying anaphylaxis.

The miR-29a has been described as a key regulator within the cardiovascular system^45^, as it is essential for the proper development and maintenance of EC functions^30^. Dysregulations of this miRNA have been associated with vascular pathological processes, such as atherosclerosis^46^, highlighting its dual role in vascular physiology and pathology. Given that cardiovascular involvement is a crucial factor in the severity of anaphylaxis^7^, it is particularly relevant to understanding the behavior of miR-29a in the endothelium. Our findings demonstrate that intracellular levels of miR-29a-3p exhibit a differential response to anaphylactic stimuli, increasing in HAECs and decreasing in HMVEC-D. The vascular tree comprises different compartments, which differ in structure and function^47^. In the context of anaphylaxis, these components respond differently to stimuli, with capillary segments and veins mainly associated with leakage and vasodilation and arteries with vasomotor tone^7^. Therefore, this contrasting intracellular pattern suggests that miR-29a-3p may exert vessel-specific effects during anaphylactic reaction, contributing to distinct adaptive or pathological mechanisms throughout the vascular system.

To evaluate the functional and molecular effects of miR-29a-3p in ECs, we performed an *in silico* analysis using its predicted target genes. Our data demonstrated that this miRNA is closely associated with extracellular matrix organization. Notably, miR-29a-3p has previously been linked to collagen alterations^39^, and the remodeling of the extracellular matrix on cardiac tissue^48^. Moreover, this miRNA has been shown to reduce inflammation and maintain extracellular matrix stability in rat models of osteoarthritis^49^. Based on all these findings, we investigate the role of miR-29a-3p in the regulation of eGCX, a structure whose integrity is compromised during anaphylaxis^13^. Inhibition of miR-29a-3p resulted in a significant reduction in *ESM1* expression. This gene encodes Endocan-1, a protein implicated in endothelial dysfunction, and whose concentration has recently been found to be elevated in plasma from a murine model of active systemic anaphylaxis^13^. Interestingly, our results suggest that miR-29a-3p can modulate *ESM1* levels in ECs highlighting the miR-29a-3p as a key regulator of eGCX dynamics and endothelial stability.

Collectively, these results highlight the dual relevance of miR-29a-3p as a promising biomarker for endotyping anaphylactic reactions, as well as a mechanistic regulator of endothelial barrier integrity.

## Authors contributions

Conceptualization: ENB and VE. Experimental design: ENB and VE. Sample collection and clinical characterization: AGL, PRR and JJL. Experimental execution and analysis: AMC, AMV, AH with participation of LPG, SFB, IdMC, ABM and ADG. Data analysis: AMC, AMV, ABM and ENB. Data interpretation: AMC, ENB and VE. Manuscript writing: AMC and AMV. Manuscript review and editing: ENB and VE. All authors contributed to the critical revision of the manuscript and approved its final version.

## Acknowledgements

This research was supported by “Instituto de Salud Carlos III” (ISCIII; PI18/00348, PI21/00158 and PI24/00360), cofounded by FEDER “Investing in your future” for the thematic network and cooperative research centers ARADyAL (RD16/0006/0013, RD16/0006/0033) and RICORS “Red de Enfermedades Inflamatorias (REI)” (RD24/0007/0018 and RD24/0007/0037). Additional support was provided by the Alfonso X El Sabio University (UAX) Foundation and the Spanish Society of Allergy and Clinical Immunology (SEAIC) Foundation. Part of the experiments for this study were carried out under an EMBO Scientific Exchange Grant (number 9328). ABM is supported by a Miguel Servet contract from the ISCIII (CP23/00046), co-funded by the European Union. SFB and ENB received a predoctoral grant (FI22/00046) and a Sara Borrell contract (CD23/00125) from the ISCIII, respectively. ENB is supported by research grants from the IIS-Princesa (PIM-011) and the UAX Foundation (950.690). LPG and IdMC are recipients of predoctoral grants (CAM_PIPF-2022/SAL-GL-24982) and Conchita Rabago Foundation contracts, respectively.

## Data availability statement

The data that support the findings of this study are available from the corresponding author upon reasonable request.

