## Supplementary material for "Circulating miR-29a as a new biomarker of food anaphylaxis and endothelial glycocalyx regulation": Sup

### Supplementary Methods

#### Ethics Committee

Authors adhered to the declaration of Helsinki and all patients were always included after giving informed consent by the donor or their relatives. Inclusion criteria were acceptance to participate in the study and an objective diagnosis of anaphylaxis. Exclusion criteria were the presence of a blood-borne disease and/or any psychic or psychological disorder that would alter the acceptance of the study.

#### Subjects of study

Patients with anaphylaxis by distinct triggers were recruited from different Spanish hospitals: Fundación Jiménez Díaz University Hospital, Central de la Cruz Roja Hospital, Guadalajara´s University Hospital, Navarra´s University Hospital, and Niño Jesús University Children´s Hospital.

#### Sample collection

All samples were obtained from patients who developed anaphylaxis accidentally. If the reaction occurred outside the hospital, acute phases were collected in emergency departments. However, if the anaphylaxis was unintentionally provoked by a controlled challenge test, samples were obtained in allergy units.

For serum processing, tubes with a separator gel (BD Vacutainer) were used. Once the peripheral blood was obtained, it was centrifuged at 1,200 g for 10 minutes at 4°C. Afterwards, samples were aliquoted in 0.5 ml and stored at -80°C until their use.

#### Circulating serum miRNAs analysis

For RNA extraction (miRNeasy Serum/Plasma Advanced kit; Qiagen), 60 µl of lysis buffer and 1 µl of UniSp2/4/5 mix (miRCURY LNA RNA Spike-in kit, Qiagen) were added to 200 µl of serum. These synthetic miRNAs serve as a quality control of the technique. Samples were incubated for 3 minutes at room temperature (RT), 30 µl of the precipitation buffer were added, and a new 3-minute incubation at RT was performed. After this, tubes were centrifuged at 12,000 g for 3 minutes, supernatants were mixed with an equal volume of isopropanol and transferred to a specific column where RNA was retained. Subsequently, three washes were performed and 15 µl of RNAase-free water was added to the column, which was incubated for 1 minute at RT and centrifuged for 1 minute to elute the isolated RNA.

Total RNA was retrotranscribed using the miRCURY LNA RT kit (Qiagen). For this purpose, 2 µl of the reaction-specific buffer, 4.5 µl of RNase-free water, 1 µl of the enzyme mixture, 2 µl of the RNA previously obtained in the extraction and 0.5 µl of UniSp6, a synthetic miRNA used as a quality control of the reverse transcription, were included for each sample. Once the mix was done, samples were placed in the PTC-100 Thermal Cycler (MJ research, Inc) where reverse transcription was carried out incubating: 60 minutes at 40°C, 5 minutes at 95°C and 5 minutes at 4°C. Finally, the cDNA obtained was stored at 20°C until further use.

For miRNAs quantification (miRCURY LNA SYBR Green PCR kit; Qiagen), 3 µl of diluted sample (1:30), 5 µl of the miRCURY SYBR Green 2 master mix, 1 µl of RNase-free water, and 1 µl of the primer of interest were added for each condition. Then, the plate was centrifuged at 1,200 g for 2 minutes at 4°C and analyzed following the next protocol: 2 minutes at 95°C; 45 cycles of amplification in two steps (first one at 56°C for 60 s and second one at 95°C for 10 s), where fluorescence was measured; a melt curve analysis at 95°C for 5 s, 60°C for 60 s and 95 °C for 1 s to inactivate the reaction; and a cooling final phase at 37°C for 30 s to cool the plate.

**Determination of miRNAs within extracellular vesicles**

For the purification of EVs (miRCURY Exosome isolation Kit – Serum and Plasma; Qiagen), 500 µl of serum were centrifuged at 12,000 g for 20 minutes at 10°C. Subsequently, 200 µl of the precipitate buffer were added and incubated for 3 hours at 4°C. Then, samples were centrifuged at 1,500 g for 30 minutes and supernatants were removed. Finally, EVs were collected adding 270 µl of the resuspension buffer.

The extraction, reverse transcription, and quantification of miRNAs were performed as previously described for circulating serum miRNA, with slight modifications. Specifically, 300 µl of serum were used, and 90 µl of lysis buffer and 30 µl of precipitation buffer were added. For qPCR, cDNA samples were diluted at 1:8.

**Western blot**

Three bona fide markers (CD9, CD63 and TSG101) of EVs were characterized by Western blot (WB). For this purpose, EVs proteins were separated by electrophoresis on sodium dodecyl sulfate polyacrylamide gels (SDS-PAGE) at 12.5% using the PowerPac Basic Power Supply section (BioRad). Subsequently, proteins were transferred to a nitrocellulose membrane (BioRad) using the Trans-Blot Turbo Transfer system (BioRad). Non-specific membrane junctions were blocked by applying a solution of phosphate buffered saline (PBS) with 0.1% Tween (Sigma) and 5% milk. Then, the membrane was incubated overnight with the corresponding primary antibodies diluted 1:500 in all cases: anti-CD9 (Invitrogen), anti-CD63 (Invitrogen), and anti-TSG101 (Abcam). For signal detection, the membrane was incubated for 1 hour with a rabbit anti-mouse (RAM) peroxidase-conjugated secondary antibody (Jackson Laboratory) diluted at 1:5,000 in 0.05% PBS-Tween solution with 3% milk. Lastly, proteins were detected in the Amersham Imager 600 system (GE Healthcare) using the chemiluminescent ECL Prime Western Blotting Detection Reagent (Thermo Scientific).

**Electron microscopy**

For the EVs visualization by electron microscopy (EM), 5 µl of the previously purified particles were fixed for 24 hours at 4°C using 50 µl of PBS with 2% paraformaldehyde (PFA, Sigma). After this, EVs were adsorbed on copper/carbon grids (Fedelco) for 3 minutes, washed with water for 1 minute, and counterstained with 1% uranyl acetate for 30 seconds. Then, samples were observed using a JEOL 1010 transmission EM (JEOL) employing a voltage of 100 kV. The analysis and image processing were carried out with the Soft Imaging Viewer software.

**Nanoparticle tracking analysis**

Nanoparticle tracking analysis (NTA) was performed at the Spanish National Cancer Research Center (CNIO, Madrid) with the NS500 nanoparticle characterization system (NanoSight) equipped with a blue laser (405 nm) and diluting samples 1:500.

**System biology analysis**

Signaling pathways analysis was performed using the ShinyGo web tool to obtain the 20 most significant pathways ranked by enrichment. Pathways within the specified size limits (minimum 2 and maximum 5000) were considered. Pathways were filtered by a False Discovery ratio (FDR) threshold of 0.05. In addition, "Remove Redundancies" option was used to eliminate pathways that share 95% of their genes and 50% of the words in their names.

On the other hand, STRING web tool was conducted to obtain information about molecular functions and biological processes. The network criteria were a minimum strength of 0.01 and a minimum of 2 counts in the network.

**Human endothelial cells culture**

HAEC and HMVEC-D were maintained in EGMTM-2 and EGM-2MV BulletKit medium (Lonza), respectively. These media were supplemented with 20% FBS, 10 µl/ml heparin and 5 µl/ml endothelial cell growth factor (ECGF). Media was changed every 3-4 days.

**Determination of miR-29a levels in stimulated ECs**

Cells were lysed (Master Pure Complete DNA & RNA Purification Kit; Lucigen) by scrapping them after adding 300 µl of lysis buffer and 1 µl of Proteinase K. The lysate obtained was collected and incubated at 65°C for 15 minutes. Then, samples were incubated at 4°C for 5 minutes and 150 µl of precipitation buffer were added. Subsequently, each sample was centrifuged at 10,000 g for 10 minutes and supernatants were collected. An equal volume of isopropanol was added, and mixtures were centrifuged again. Next, to remove the remaining DNA, the precipitate was resuspended in 200 µl of DNA nuclease buffer and 5 µl of DNA nuclease I. Consecutively, samples were incubated at 37°C for 30 minutes and 200 µl of both the lysis and precipitation buffers were added. Then, samples were incubated at 4°C for 5 minutes and centrifuged at 10,000 g for 10 minutes at 4°C. After this, an equal volume of isopropanol was added, and samples were centrifuged again. Finally, the RNA extracted was resuspended in 20 ml of trisaminomethane buffer (TE) and 1 µl of RNAse inhibitor. Finally, reverse transcription and miRNAs quantification were performed as described in the circulating serum miRNAs section.

**Transfection of miRNA and gene expression analysis in HAEC**

For RNA extraction, cell scrapers and 1 ml of TRIzol (Invitrogen) were employed to lysate cells. This suspension was subjected to a chloroform gradient (200 µl) by centrifugation at 14,000 rpm and 4°C for 15 minutes for RNA obtaining. Then, RNA was mixed with 500 µl of isopropanol and incubated overnight at 4°C to facilitate RNA precipitation. After this, it was centrifuged at 14,000 rpm and 4°C for 15 minutes, and the supernatant was discarded. To clean the RNA, two additional centrifugations were performed at maximum speed using 70% and 100% ethanol. Finally, the RNA was resuspended in 10 µl of RNase-free water, and its concentration was measured in a NanoDrop (ND1000).

### Supplementary Figures

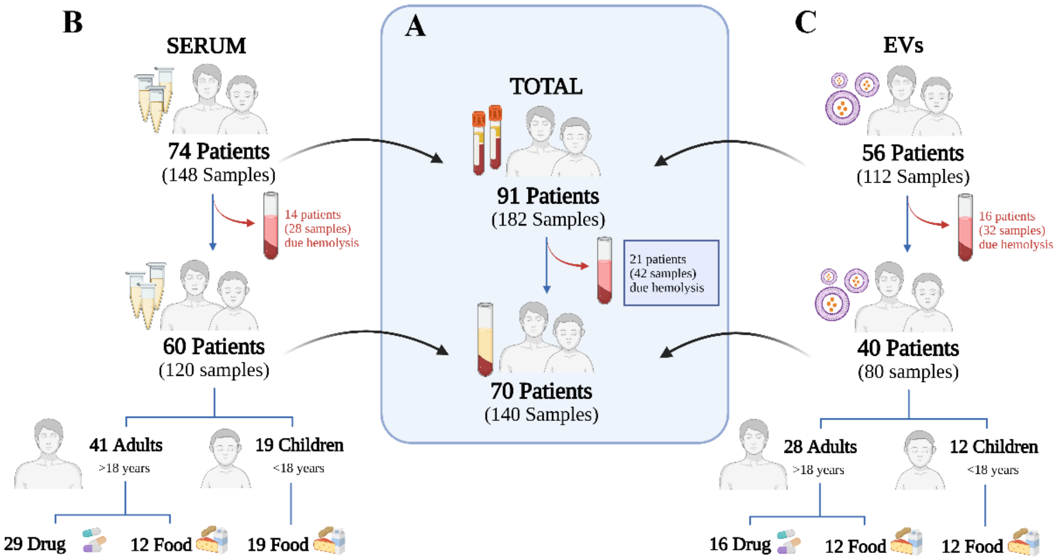


**Supplementary Figure S1. Diagram of patient distribution used in this study.** (**A**) The initial patient cohort included 91 patients with anaphylaxis by different triggers. However, duo to the haemolysis in one, or both, of their samples, 21 patients were discarded, resulting in a final cohort of 70 patients with anaphylaxis (**B**) For serum studies, a total of 74 patients with anaphylaxis were included. However, 14 of them were excluded due to haemolysis in one, or both, of their samples, resulting in a final cohort of 60 patients (41 adults and 19 children). Among adults, 29 were AD and 12 AF. (**C**) For the study of miRNAs within extracellular vesicles (EVs), a total of 56 patients with anaphylaxis were included. However, 16 of them were discarded due to haemolysis in one, or both, of their samples, giving rise to a final cohort of 40 patients (28 adults and 12 children). Among adults, 16 were AD and 12 were AF.

### Supplementary Tables

**Supplementary Table 1**. Clinical characteristics of patients and features of their hypersensitivity reactions detailed per subject.

| | | Patients | | | | Clinical symptoms | | | | | | Severity | Treatment (before collecting acute sample) | | | | | Tryptase (ng/mL) | | Use | |
|---|---|---|---|---|---|---|---|---|---|---|---|---|---|---|---|---|---|---|---|---|---|
| | | Number | Age | Gender | Trigger | Skin | Mucous | Digestive | Respiratory | Nervous | Vascular | Grade | Epinephrine | H1R antagonist | H2R antagonist | Corticosteroids | β2-adrenergic agonist | Acute | Basal | Serum | EVs |
| ADULTS | DRUGS | 1 | 44 | F | AAS | Er; Pr | | Dp; Ns; Ap | Dy; CT; Cg | | | 2 | | X | | X | | 4.2 | 6.07 | X | X |
| | | 2 | 43 | F | Ibuprofen | Er; Pr | | | Dy; Sz; NC | | | 2 | X | X | | X | | 3.48 | 3.58 | X | |
| | | 3 | 37 | F | Amoxicillin/Clavulanic acid | Er; Pr | Cj | Ns; Ap | Dy; Sz | | | 2 | X | | | | | 5.39 | 5.4 | X | |
| | | 4 | 65 | F | Amoxicillin | Pr; Ur | | Ns; Ap; Vm; Dr | Wz | Dz | | 3 | | | | | | 6.94 | 5.37 | X | X |
| | | 5 | 43 | F | Vancomycin | Er; Pr | | Dp | | | | 2 | | X | | X | | 3.67 | 3.86 | X | |
| | | 6 | 31 | F | Taxol | Er | | Ns; Ap | Dy; Hp | | | 3 | X | X | | X | | 3.92 | 5.43 | X | X |
| | | 7 | 49 | F | Amoxicillin/Clavulanic acid | Er; Pr | Ag | Ns; Ap | Wz; Cg | | | 2 | X | | | X | | 5.32 | 5.82 | X | X |
| | | 8 | 29 | F | Codeine | Er; Pr; Ur | | | Wz; Cg | | Hy | 3 | X | | | | | 3.06 | 3.36 | X | X |
| | | 9 | 50 | M | Augmentin | Er; Pr; Ur | Ag | Ns; Ap | Dy; Hp | Dz | Tc; Hy | 3 | X | X | | X | | 12 | 8.63 | X | |
| | | 10 | 62 | M | Amoxicillin/Clavulanic acid | Er; Pr; Ur; Hb | Cj | | Sz; Rh | | | 2 | | | | X | | 3.4 | 3.95 | X | |
| | | 11 | 44 | F | Ciprofloxacin | Er | Ag | Ns; Ap | Sz; Rh; NC | | | 2 | X | X | | X | | 3.64 | 4.46 | X | X |
| | | 12 | 56 | F | Levofloxacin | Er; Pr | | | Dy; Hp | | Tc | 3 | X | X | X | X | | 7.6 | 6.16 | X | |
| | | 13 | 49 | F | Celecoxib | Er; Pr | | | Sz; NC; Cg | | | 2 | | X | | X | X | 3.12 | 3.69 | X | |
| | | 14 | 73 | M | Contrast | Pr | | | Wz | Dz; Sw | Sy | 3 | | X | X | X | | 13 | 4.38 | X | |
| | | 15 | 22 | M | Contrast | Er | Ag; Cj | | Sz; Rh | | | 2 | X | X | | X | | 28.6 | 4.94 | X | |
| | | 16 | 22 | F | Penicillin | Ur | | PFB | Dy; Sz; Cg | | Tc | 3 | X | X | | | | 5.56 | 4.49 | | X |
| | | 17 | 40 | M | AAS | Er; Pr | | Ns; Ap | Wz; Sz; Rh | Sw | | 3 | | | | | | 5.33 | 5.05 | X | X |
| | | 18 | 49 | F | AAS | | Cj | | CT; Rh | | | 2 | X | X | | X | | 3.21 | 4.32 | X | |
| | | 19 | 22 | F | AAS | Er | Ag | | Dy | | | 2 | | | | | | 2.9 | 2.22 | X | |
| | | 20 | 36 | F | AAS | Er | Ag | Ns; Ap | Sz; Rh | | | 2 | X | | | | | 5.24 | 7.47 | X | X |
| | | 21 | 50 | M | Ibuprofen | Er; Pr | Ag | | Cg | | | 2 | | | | | | 4.28 | 4.45 | X | |
| | | 22 | 47 | F | Amoxicillin/Clavulanic acid | E; Pr; Ur | | Ns; Ap | Sz | Dz | Hy | 3 | X | | | | | 15.2 | 8.48 | X | |
| | | 23 | 76 | F | Metamizole | Er; Pr | | Ns; Ap | Wz | | Hy | 3 | | X | | X | X | 48 | 7.83 | | X |
| | | 24 | 33 | M | Metamizole | | | Ns; Ap; Vm; Dr | Bc; Wz | | | 2 | X | X | | X | X | 5.34 | 4.38 | X | |
| | | 25 | 37 | F | Penicillin | Er | | Ns; Ap | Dy; CT; Cg | Dz | Tc | 3 | X | | | | | 10.8 | 7.35 | X | X |
| | | 26 | 59 | F | Amoxicillin | Er | | Ns; Ap | | Dz | Tc; Hy | 3 | X | | | | | 28.4 | 8.1 | | X |
| | | 27 | 56 | F | AAS | Er; Pr | | | | Dz | Hy; Sy | 3 | X | X | | X | | 3.98 | 3.09 | X | |
| | | 28 | 43 | F | Taxane | | | Ns; Ap | Dy | Dz | | 3 | | X | | X | X | 2.24 | 1.5 | X | |
| | | 29 | 67 | M | Taxane | | | | Dy; CT; Hp | GD | | 3 | X | | | X | | 7.77 | 5.48 | X | |
| | | 30 | 38 | F | Amoxicillin/Clavulanic acid | Er; Pr | Ag | Ap | Dy; CT | | Hy | 3 | X | | | X | | 6.63 | 3.46 | X | |
| | | 31 | 69 | M | Oxaliplatin | Er | Ag | | Bc; Sz; Hp | | Tc; Hy; Sy | 3 | X | X | | X | X | 45.5 | 4.68 | | X |
| | | 32 | 27 | M | AAS | Ur | Ag | | Dy; CT | | | 2 | | | | | | 2.83 | 2.72 | X | X |
| | | 33 | 69 | M | Contrast | Er; Pr | | | Dy; Wz | Dz | Tc; Hy; Sy | 3 | X | X | X | X | | 106 | 4.33 | X | X |
| | | 34 | 42 | M | Diclofenac | Pr; Ur | Ag | PFB | Rh; NC; Cg | | | 2 | X | X | | X | | 6.84 | 4.58 | | X |
| | FOOD | 35 | 59 | M | Anchovy | Er; Pr | Ag | Vm | Bc | | | 2 | | X | X | X | X | 7.62 | 4.41 | X | |
| | | 36 | 20 | F | Nuts | | | Vm | Bc; Rh; Hp | | Hy | 3 | | | | X | | 17.6 | 6.18 | X | |
| | | 37 | 39 | F | Emperor | Er; Pr; Ur | | Ns; Ap; Vm; Dr | | | | 2 | | | | | | 8.64 | 7.41 | X | X |
| | | 38 | 33 | F | Nuts | Er; Pr; Ur | Ag | | Sz; Rh; Cg | | | 2 | X | X | X | X | | 12.6 | 7.2 | X | |
| | | 39 | 29 | M | Mustard | Pr | Ag | Ns; Ap; Vm; Dr | Sz; NC | | | 2 | X | | | | | 6.29 | 4.17 | | X |
| | | 40 | 45 | M | Honey | Ur | Ag | Ns; Ap | Wz | | | 2 | X | X | X | X | | 18.4 | 6.87 | X | X |
| | | 41 | 36 | F | Peach | Er; Pr; Ur | Ag | Ns; Ap | Dy; NC | | Tc | 2 | X | | | | | 4.34 | 3.75 | X | X |
| | | 42 | 47 | M | Honey | Pr; Ur | Ag | Ns; Ap | | | Sy | 3 | X | X | | X | | 9.36 | 3.07 | | X |
| | | 43 | 21 | F | Lettuce | Ur | Ag | | Dy | | | 2 | | X | | X | | 3.53 | 1.33 | X | |
| | | 44 | 28 | F | Paraguayan | Pr | Ag; Cj | Ns | | | Hy | 3 | X | X | | X | | 3.48 | 1.65 | X | |
| | | 45 | 43 | F | Prawns | Ur | Ag | Ns; Ap; Vm; Dr | | | Hy; Sy | 3 | X | X | X | X | | 31.7 | 6.44 | | X |
| | | 46 | 38 | M | Peach | Pr; Ur | | Ns; Ap; Dr | Sz | | | 2 | | X | | X | | 13.3 | 6.14 | X | X |
| | | 47 | 24 | M | Squid | Er; Pr | | Ns; Ap; Vm; Dr | Sz; NC | | | 2 | | X | | X | | 13 | 7.25 | X | X |
| | | 48 | 20 | F | Nuts | Er | Ag | | Dy; Wz; Hp | | | 3 | | | | | | 7.35 | 4.06 | | X |
| | | 49 | 30 | F | Kiwi | Er; Pr; Ur | Ag | Ns; Ap; Vm; Dr | Wz; Sz | | | 2 | | X | | X | | 14.2 | 3.65 | X | X |
| | | 50 | 19 | M | Cheese | E; Pr | Ag; Cj | | Rh; NC; Dy | | | 2 | X | X | | | | 14 | 5.15 | X | X |
| | | 51 | 68 | F | Fish | Pr | | Ap | Dy; Wz; CT; Hp | | | 3 | X | X | | X | X | 14.3 | 7.27 | | X |
| CHILDREN | FOOD | 52 | 17 | F | Milk | Er; Pr | | Ns; Ap | | | | 2 | | | | | | 6.93 | 5.91 | X | X |
| | | 53 | 11 | F | Egg | | | Ns; Ap; Vm; Dr | Sz | | | 2 | X | | | | | 5.83 | 3.34 | X | X |
| | | 54 | 4 | M | Cashew | Er | Ag | | Sz | | Tc | 2 | | | | | | <1 | <1 | X | X |
| | | 55 | 5 | F | Rooster | Er; Pr; Ur | | | Wz | | | 2 | | X | | | X | 9.36 | 11.4 | X | |
| | | 56 | 9 | F | Milk | | | Ns; Ap; Vm; Dr | Sz | | | 2 | X | | | | | 5.5 | <1 | X | |
| | | 57 | 9 | M | Egg | Er; Pr | | Ns; Ap; Vm; Dr | Sz | | Tc | 2 | | | | | | 5.21 | 3.01 | X | X |
| | | 58 | 14 | F | Egg | Er; Pr | Ag | Ns; Ap | Sz | Dz | Hy | 3 | | | | | | 2.6 | 2.29 | X | X |
| | | 59 | 11 | F | Milk | Ur | Ag | | Wz; Sz | | | 2 | | | | | | 3.81 | 3.31 | X | X |
| | | 60 | 15 | M | Cheese | Er | | | Wz; Sz | | | 2 | | | | | | 8.03 | 5.69 | X | X |
| | | 61 | 13 | F | Milk | | | Ns; Ap | Wz; Sz | | | 2 | | | | | | 5.04 | 2.98 | X | X |
| | | 62 | 17 | M | Hazelnut | | Ag | | Dy | | | 2 | X | X | | | | 5.62 | 7.04 | X | X |
| | | 63 | 5 | M | Milk | Er | | Ap; Vm; Dr | Dy | | Hy | 3 | X | X | | X | X | 7.83 | 4.97 | X | X |
| | | 64 | 14 | M | Egg | Er | Ag | Ns; Ap; Vm; Dr | Sz | | | 2 | | | | | | 2.88 | 2.44 | X | X |
| | | 65 | 10 | F | Egg | Er; Pr | Ag | | Sz | | | 2 | | | | | | 3.89 | 3.78 | X | |
| | | 66 | 14 | F | Nut | Ur | Ag | | Cg | Dz | | 3 | X | X | | X | | 6.63 | 5.65 | X | X |
| | | 67 | 7 | M | Rooster | | Ag | | Cg | | | 2 | X | X | | X | | 3.84 | 2.53 | X | |
| | | 68 | 14 | F | Milk | Ur | | Ns; Ap | Wz; Sz; Rh; NC | | | 2 | X | X | | X | X | 8.74 | 1.72 | X | |
| | | 69 | 11 | F | Milk | | | Ns; Ap | Sz; Cg | | | 2 | X | X | | | X | 8.57 | 7.05 | X | |
| | | 70 | 15 | M | Peanut | Ur | Ag | PFB | Dy | | | 2 | X | | | X | | 3.03 | 2.62 | X | |

Gender: M (male), F (female). Clinical symptoms—skin: Er (erythema), Ur (urticaria) and Pr (pruritus); mucous: Ag (angioedema) and Cj (conjunctivitis); digestive: Ns (nausea), Vm (vomit), Dr (diarrhea), Ap (abdominal pain), Dp (dysphagia) and PFB (pharyngeal foreign body); respiratory: Dy (dyspnea), Wz (wheezing), Rh (rhinitis), Cg (cough), Sz (sneeze), Bc (bronchospasm), CT (chest tightness), NC (nasal congestion) and Hp (Hypoxia); neurological: Dz (dizziness), Sw (sweating) and GD (general discomfort); vascular: Hy (hypotension), Tc (tachycardia) and Sy (syncope).

**Supplementary Table 2**. Primers used for miRNAs analysis.

| Primers | Sequences | References |
|---|---|---|
| **hsa-miR-451a** | 5’ AAACCGUUACCAUUACUGAGUU 3’ | *YP02119305* |
| **hsa-miR-23a-3p** | 5’ AUCACAUUGCCAGGGAUUUCC 3’ | *YP00204772* |
| **hsa-miR-30e-5p** | 5’ UGUAAACAUCCUUGACUGGAAG 3’ | *YP00204714* |
| **hsa-miR-29a-3p** | 5’ UAGCACCAUCUGAAAUCGGUUA 3’ | *YP00204698* |
| **hsa-miR-29a-5p** | 5’ ACUGAUUUCUUUUGGUGUUCAG 3’ | *YP00204430* |

**Supplementary Table 3.** Primers used to detect endothelial glycocalyx (eGCX) mRNA in human aortic endothelial cells (HAEC). Fibronectin 1 (*FN1*), Rho-associated protein kinase 1 (*ROCK1*), Claudin 5 (*CLDN5*), Rac family small GTPase 1 (*RAC1*), Fibrillin 1 (*FBN1*), Endocan 1 (*ESM1*), Syndecan 1 (*SDC*), Aggrecan (*ACAN*), Heparan sulfate 2-O-sulfotransferase 1 (*HS2ST1*), N-Deacetylase and N-Sulfotransferase 2 (*NDST2*).

| Genes | Primers | Sequences |
|---|---|---|
| ***FN1*** | Forward | 5’ CGAGGTGACAGAGACCACAA 3’ |
| | Reverse | 5’ CTGGAGTCAAGCCAGACACA 3’ |
| ***ROCK1*** | Forward | 5' GACTGGGGACAGTTTTGAGAC 3' |
| | Reverse | 5' GGGCATCCAATCCATCCAGC 3' |
| ***CLDN5*** | Forward | 5' CCTATGACCCCAGTCAATGC 3' |
| | Reverse | 5' GGGAAATACGTGTGA 3' |
| ***RAC1*** | Forward | 5' GAGCAGAAGCTGATCTCCGAGGAG3' |
| | Reverse | 5' TTACAACAGCAGGCATTTTCTCTT 3' |
| ***FBN1*** | Forward | 5' GGASTGACATCAGCAGGCAC 3' |
| | Reverse | 5' TACACAAATCCTTTGGGGCA 3' |
| ***ESM1*** | Forward | 5’ CTGGAGCGCCAAATATGCG 3’ |
| | Reverse | 5’ TGAGACTGTACGGTAGCAGGT 3 |
| ***SDC1*** | Forward | 5’CCTTGTCAGGGTAGACAGCCT 3’ |
| | Reverse | 5’ GACAGAGGTAAAAGCAGTCTCG 3’ |
| ***ACAN*** | Forward | 5’ CGCCACTTTCATGACCGAGA 3’ |
| | Reverse | 5’ TCATTCAGACCGATCCACTGGTAG 3’ |
| ***HS2ST1*** | Forward | 5' TATGATGCCGCCCAAGTTG 3' |
| | Reverse | 5' CTGTTCAATTTCTCGGACTTCGT 3' |
| ***NDST2*** | Forward | 5' CTGCTGATTGGTTTCAGTCTT 3' |
| | Reverse | 5' CCACTGCTACTACAGTCTCCC 3' |
| ***18S*** | Forward | 5’ CCGTCGTAGTTCCGACCATAA 3’ |
| | Reverse | 5’ CAGCTTTGCAACCATACTCCC 3’ |
